# Visualization of a Standing Lattice of Scattered Light on Spherical Water Droplets under an Acoustic Field Using Biogenic Microparticles

**DOI:** 10.1101/2021.11.03.467206

**Authors:** Masakazu Iwasaka

**Affiliations:** Hiroshima University, 1-4-2 Kagamiyama, Higashihiroshima, Hiroshima 739-8527, Japan

**Keywords:** light scattering, acoustic field, microparticle, water droplet, standing wave

## Abstract

Micromanipulation using acoustic sound is a promising technique for drug delivery, cell manipulation, biosensors, and microfluidic devices. Additionally, the visualization of acoustic fields by advanced optical measurement techniques can be combined with this micromanipulation technique. The present study reveals that a lattice pattern of reflected light appears on the surface of water droplets containing microparticles when the droplets are exposed to audible sound in the range of 1900–10000 Hz. A piezoelectric membrane providing an audible acoustic field induced a stream of microparticles on which the lattice pattern overlapped, with the appearance of a standing wave. The effects of microparticles, including BaSO_4_, TiO_2_, and guanine platelets derived from fish scales, on the formation of the lattice pattern were investigated. These three types of microparticles in water enabled a visualization of the vortex streams and generated a lattice pattern of reflected light. The guanine platelets exhibited the most precise lattice pattern over the droplet surface, with a lattice width of 100–200 μm. This phenomenon may provide a new tool for detecting and manipulating micro vortex flows in the aqueous chamber of a microfluidic device combined with an acoustic transducer.

## INTRODUCTION

Numerous studies have focused on possible applications of acoustic waves for micro-to millimeter structures. For example, methods for altering the mechanical condition of microbubbles coated by lipid molecules have been investigated for improving drug delivery systems,^1^ and vibration and shrinking at the surface of microbubbles with lipid membranes have been analyzed.^2,3^

To target micromechanical conditions, various approaches have been applied for liquid phases. To explore the effect of ultrasonic waves on a liquid stream, a method for controlling a water stream inside a living cell has been investigated.^4^ The mechanism by which acoustic fields affect streams in micrometer spaces may enable flow control in microfluidic circuits. A recent work reported on a surface acoustic wave device that manipulates microparticles and cells using standing waves.^5^

The use of ultrasonic waves in microfluidic circuits arose during the early stages of microfluidic device development.^6^ Acoustic waves applied to liquid droplets and flows enable the trapping of microbeads^7^ and microalga.^8^ Furthermore, acoustic fluidic devices may contribute to environmental sustainability by collecting microplastics.^9^

Acoustic waves are also useful for manipulating liquid boundaries^10^ and particles^11^ in dynamic conditions. For example, this approach has been applied to manipulate latex particles and yeast cells. A standing wave produced by an ultrasonic wave can efficiently control the alignment of microbeads and can improve the sensitivity of quartz crystal microbalance biochips.^12^

It is anticipated that the combination of ultrasonic waves with other fields will lead to various types of methods for sensing and actuating. Numerous new technologies have been developed through the combination of ultrasonic waves with light. The use of light is advantageous for achieving high sensitivity and remote sensing availability. Whereas many applications have been developed for light waves combined with ultrasonic sound, fewer applications have been reported for light waves combined with audible sound.

Raman–Nath diffraction is a classical phenomenon in the fusional field of audible sound and visible light.^13-16^ This phenomenon is a type of light diffraction and is related to recent light technologies for sound sensing. Optical microphones and techniques for visualizing ultrasonic waves have been reported.^17-21^ It is important to explore these new innovations on the basis of acoustic manipulation and light measurement technology.

The present study explores the effects of audible sound on light-scattering patterns in water droplets containing floating microparticles. The effect on light scattering caused by biogenic guanine particles (platelets) under audible sound exposure was compared with that of barium sulfate and titanium oxide particles. Here, biogenic guanine platelets from fish scales were used, because previous studies have shown that these platelets exhibit a unique property as a liquid stream tracer.^22,23^ The platelets on the body surface of fish strongly reflect light and sometimes contribute to the formation of structural color.^24-27^ This fish platelet consists of a nucleic acid base molecule, guanine, and is very thin (~100 nm). Most of the platelet is composed of anhydrous guanine crystal.^28,29^ We previously reported that these fish guanine platelets orient parallel to applied magnetic fields and dynamically change light reflection based on the magnetic orientation when floating in water.^30,31^ In addition, these platelets exhibit a distinct light reflection anisotropy and cause strong light interference when they are aligned in parallel.^32, 33^ This work shows that guanine platelets floating in a water droplet exhibit a self-organized lattice pattern of scattered light under audible sound exposure at 1900–10000 Hz.

## EXPERIMENTAL SECTION

Figure 1 shows the experimental setup for observing the effects of acoustic field stimulation on the optical surface image of a water droplet. Incident light was provided from the side by directing a light guide connected to a white light emitting diode (LED) light source (LA-HDF158A, Hayashi Repic Co. Ltd., Japan). The microscopic observation system consisted of a high-resolution Navitar 2.0 × 1-51473 microscopic lens and a CMOS camera (Hozan L-835), as shown in Figure 1a. The white balance and exposure times were controlled manually. Figure 1b shows the side and top views of the piezoelastic transducer and a water droplet containing floating particles. Here, a 40- to 80-μL water droplet was placed on a dome-shaped frontal cover on a piezoelectric membrane plate (piezo spherical dome tweeter with flare, L001, Kemo Electronic GmbH, Geestland, Germany).

**Figure 1.**
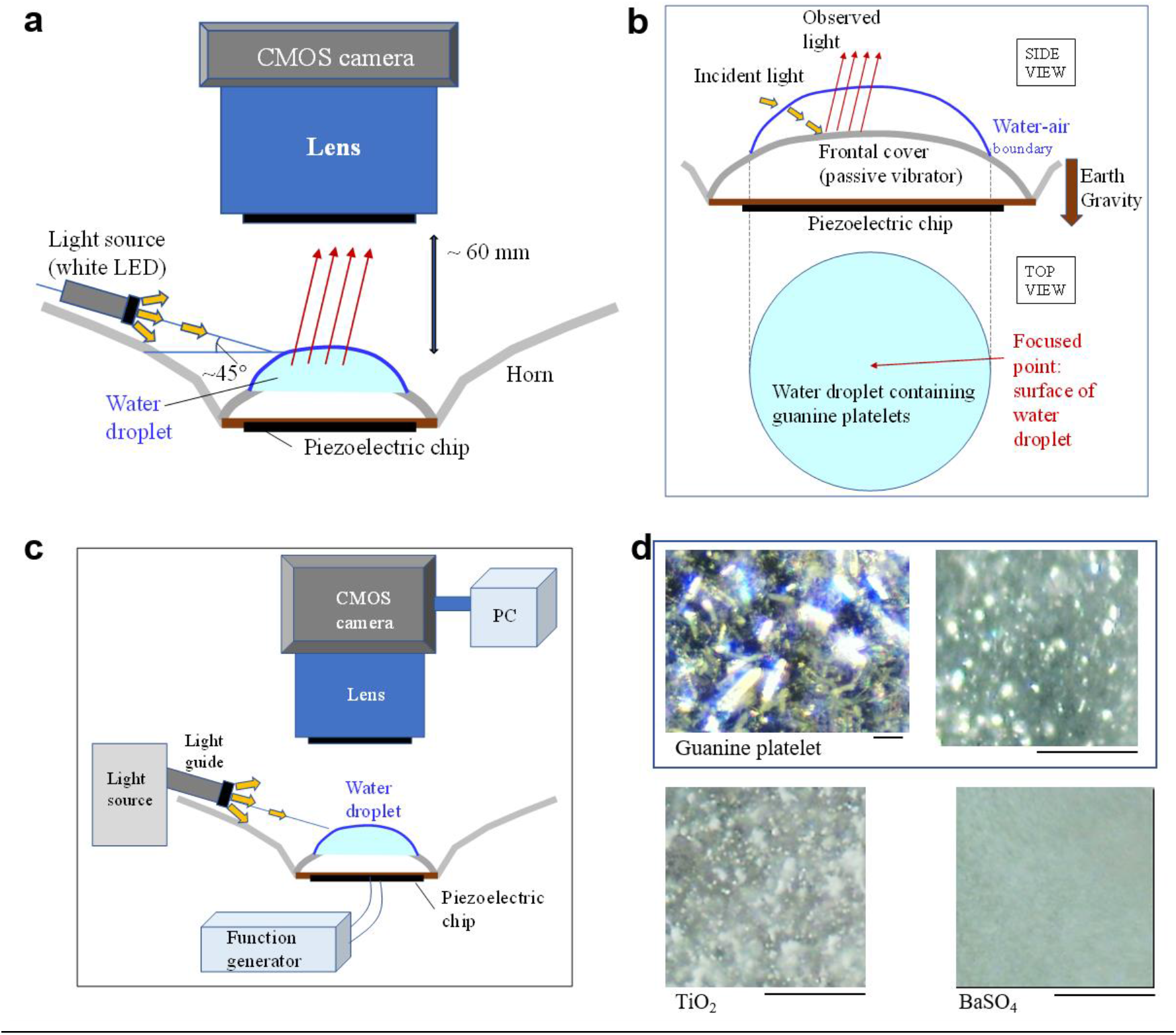
Experimental setup for observing the surface of a water droplet under acoustic field stimulation. (a) Configuration of the light source, observed sample, and microscope system. (b) Side and top views of the acoustic field transducer and water droplet containing guanine platelets, with incident light provided from the side.

To the terminals of the piezoelectric plate, a voltage was supplied as a rectangular wave form (with 50% duty) from a functional generator (Multifunction Generator WF1968, NF Corporation, Tokyo, Japan). The peak-to-peak amplitude of the supplied voltage was 20 V (i.e., +/− 10 V). The frequency of the voltage input to the piezoelectric membrane varied from 500 to 40000 Hz, and changes on the surface of the water droplet were observed to determine the specific frequency at which vibration resonance occurred.

Three types of microparticles, i.e., guanine platelets from fish, TiO_2_ particles, and BaSO_4_ particles, were used for the experiments. Biogenic guanine platelets were obtained from the scales of goldfish, *Carassius auratus*. Platelets were extracted from the scales in distilled water in a centrifugal tube (Corning, 430790, 15 mL) with a spatula (As-one, 1-9404-01, PP Spatula, 150 mm). The protocol for the fish treatment was approved by the Hiroshima University Animal Care and Use Committee (approval numbers F16–2 and F20-4, Hiroshima University). The extracted fish guanine platelets were stored with 0.01% sodium azide in distilled water for a few weeks prior to the acoustic field exposure experiments. The final concentration of the platelets in water was approximately 10^6^–10^7^ per millimeter for the acoustic field experiments. TiO_2_ powder was purchased from Benifuji (Tokyo, Lot No. 190418), and BaSO_4_ powder was obtained from Wako Pure Chemical Industries (Osaka, 022-00425). TiO_2_ and BaSO_4_ powders were also suspended in distilled water, and the supernatant of the suspension was used.

## RESULTS AND DISCUSSION

Thin guanine platelets, which typically have dimensions of 5–10 μm × 20–40 μm × 110 nm, floated in the water for 1 hour without apparent sedimentation. While the platelets floated in water, incident light supplied from the side resulted in a shining water droplet, as shown in Figure 2a. In the absence of an acoustic field, the water droplet containing guanine platelets showed a maximum reflection spot in the center of the surface, corresponding to the regular reflection obtained by the frontal cover of the piezo transducer. For the demonstration shown in Figure 2, a stream on the surface of the water droplet appeared at 1840 Hz. The stream was clearly visualized by platelets that aligned along the stream. For an applied acoustic field at 1870 Hz, a lattice or cellular pattern appeared around the spot of the regular reflection. The lattice pattern maintained a constant position even though the water stream on the droplet continued to move. Initially, it was conjectured that the lattice or cellular pattern was a standing wave produced by the applied acoustic field on the water droplet. However, as discussed in the following results, it is thought that the lattice pattern is generated by scattered light propagating inside the water droplet containing microparticles. Figure 2b shows expanded images of the scattered light on the water droplet. The light diffraction pattern on the water surface, which resembles a cellular pattern, changed its shape slightly during the observation period.

**Figure 2.**
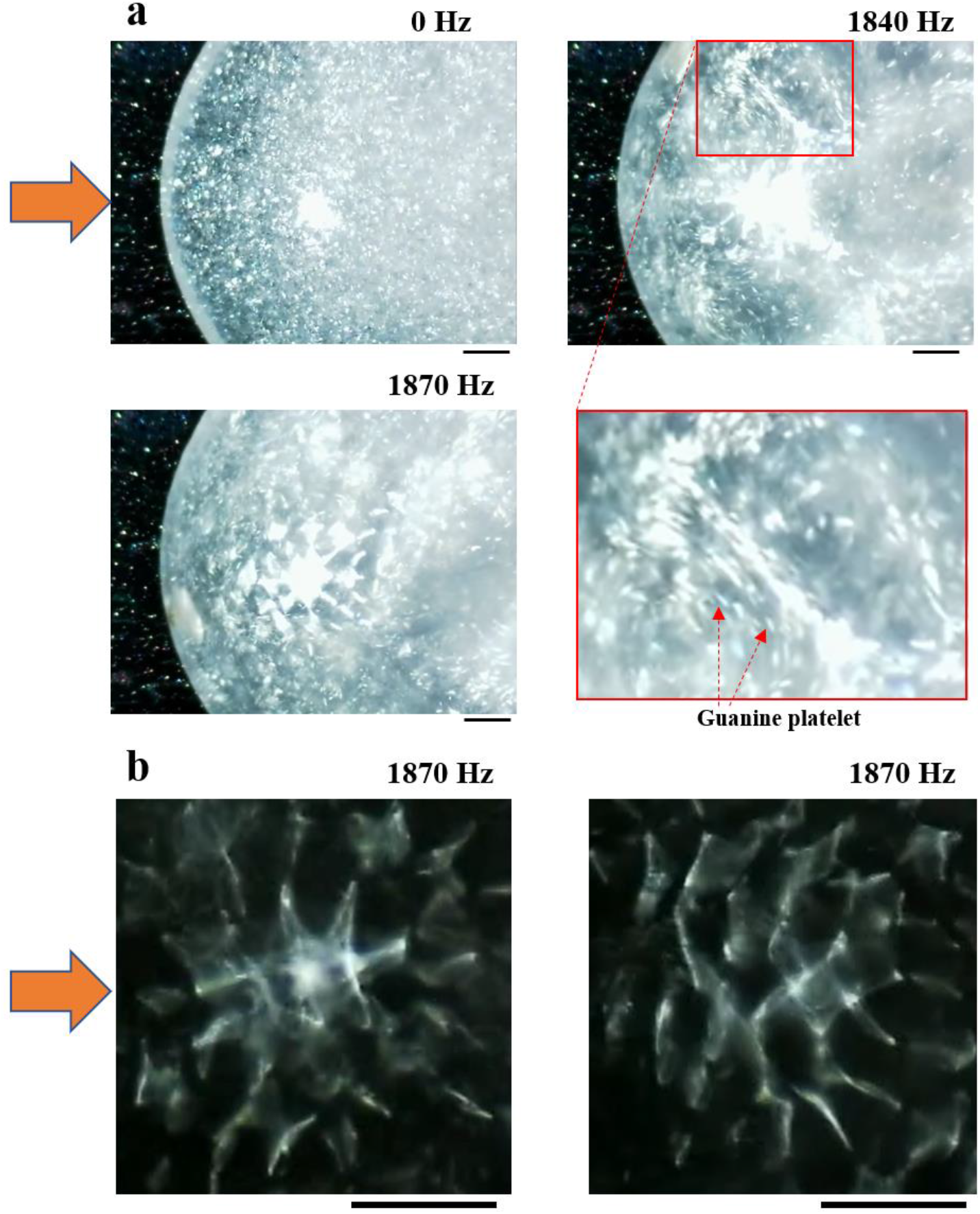
Appearance of standing lattice patterns of scattered light on a water droplet containing thin guanine microplatelets when exposed to acoustic field frequencies of up to 1870 Hz. Incident light was provided from the left-hand side. (a) Comparison of light-scattering patterns with and without acoustic field stimulation. (b) Two examples of visualized standing waves on a water droplet containing guanine platelets with acoustic field stimulation at 1870 Hz. The orange arrow indicates the direction of incident light on the horizontal plane. The scale bar represents 1 mm.

As the frequency of the rectangular wave varied, a range of frequencies showed no response in the water stream or standing patterns whereas a distinct response occurred near the resonance frequency. The resonance frequency is expected to be related to the acoustic field wave propagation inside a water droplet with a specific volume and shape because the resonance frequency varied as the sample size on the transducer cover plate changed. Figure 3 shows the standing lattice pattern of scattered light on a water droplet containing thin guanine microplatelets at higher acoustic field frequencies of 5870–7420 Hz. Here, a more precise structure of the cellular components was observed than that at 1840 Hz.

**Figure 3.**
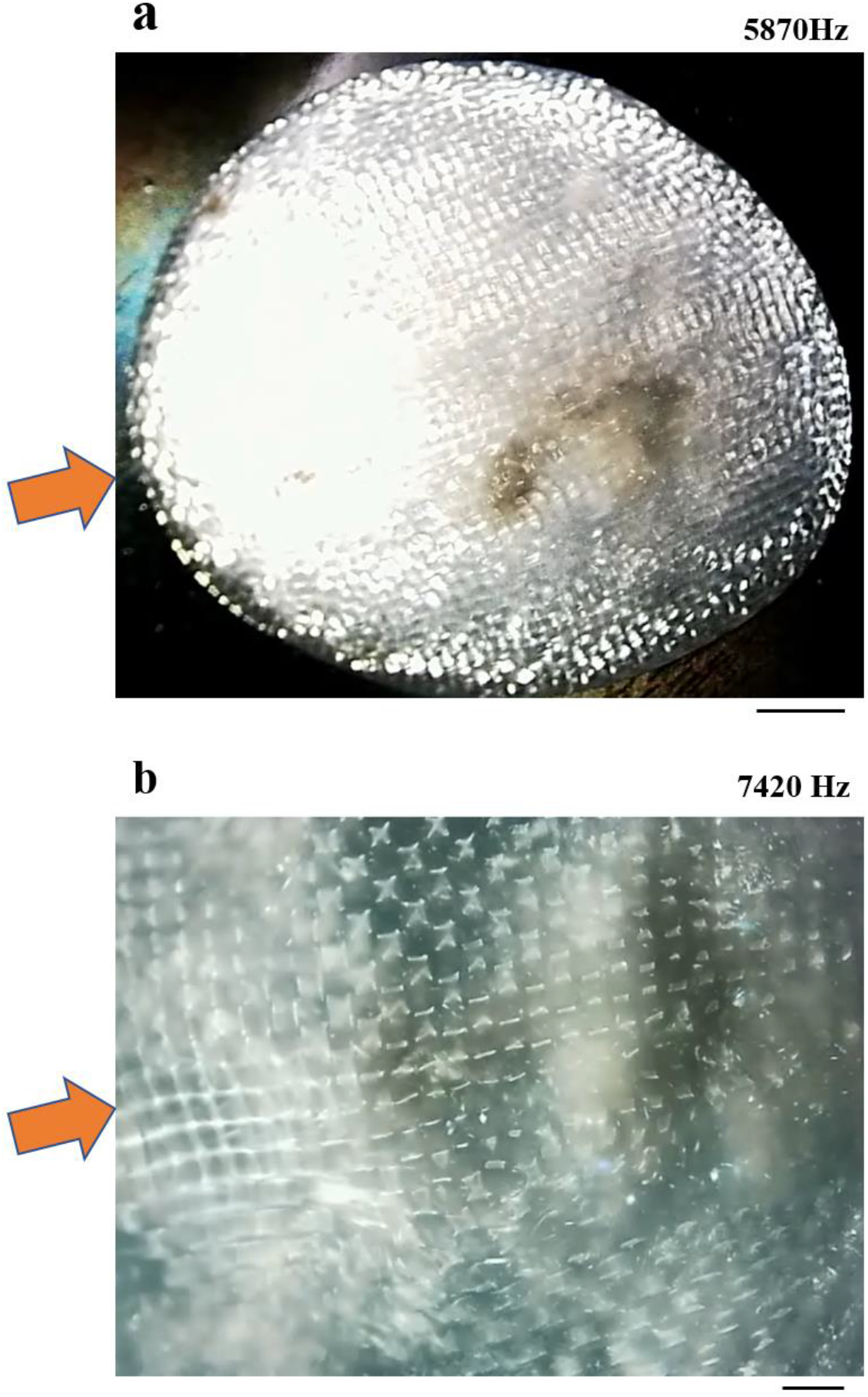
Observation of a standing lattice pattern of scattered light on a water droplet containing thin guanine microplatelets when exposed to acoustic field stimulation at (a) 5870 and (b) 7420 Hz. Incident light is presented as an orange arrow. The scale bars for (a) and (b) represent 1 mm and 300 μm, respectively.

To compare water droplets with different contents, the same experiments were performed on a pure water droplet (distilled water) and on droplets containing other types of microparticles, such as BaSO_4_ and TiO_2_ microparticles. Figure 4 shows a standing wave at the edge of the water droplet in the absence of microplatelets when an acoustic field was applied at 1900 Hz. In this case with water only, it is thought that the standing waves near the surface of the water droplet caused a cyclic gradient of the refractive index, which generated a pattern of scattered light.

**Figure 4.**
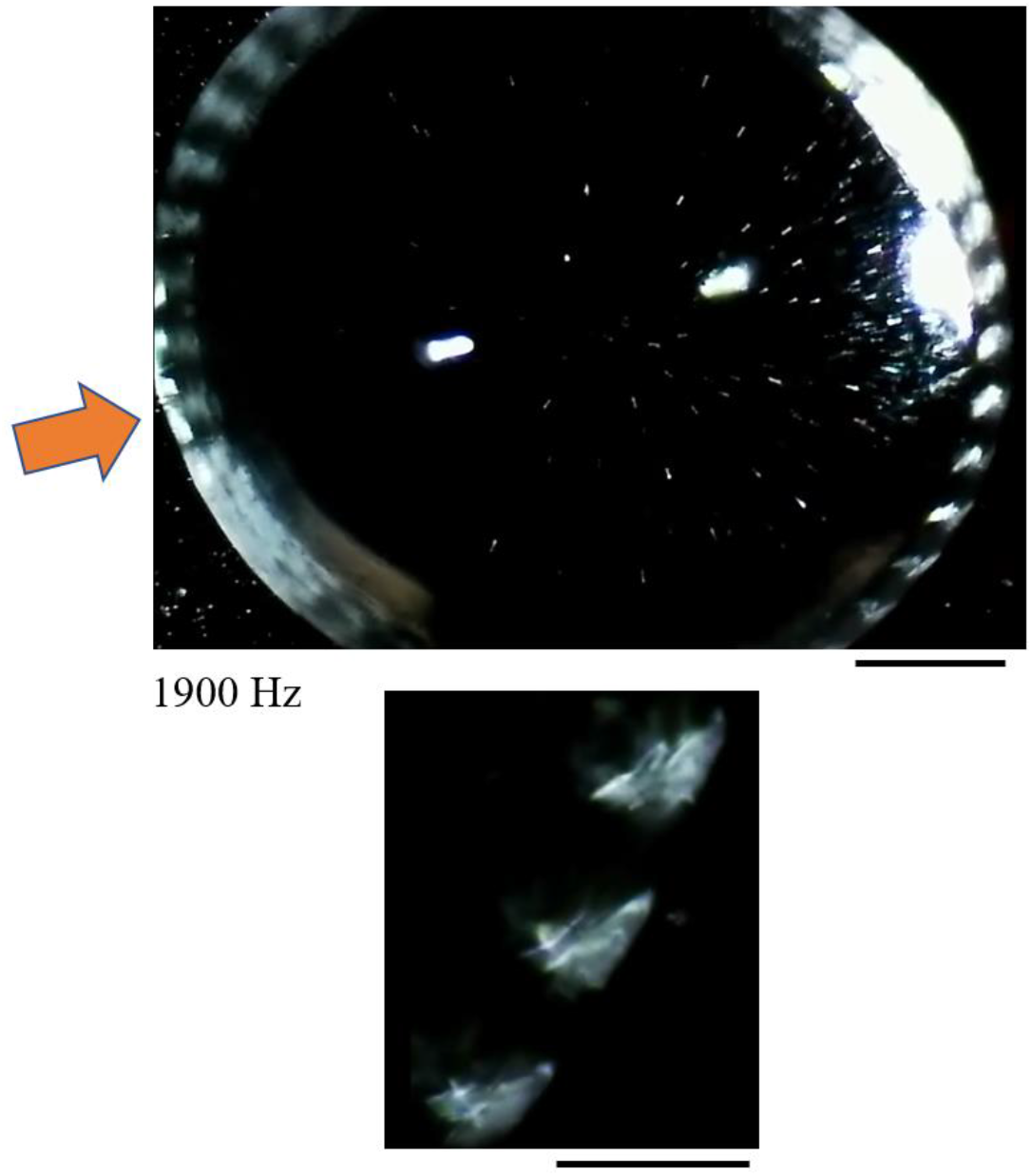
Observation of a standing wave at the edge of a water droplet without microplatelets when exposed to acoustic field stimulation at 1900 Hz. Incident light is presented as an orange arrow. The scale bars for the top and bottom panels represent 1 mm and 0.5 mm, respectively.

The effect of the type of floating microparticles in water on the standing lattice pattern of scattered light was investigated for three types of microparticles, i.e., guanine platelets from fish, TiO_2_ microparticles, and BaSO_4_ microparticles (Figure 5). The images show representative patterns obtained for each type of material at frequencies of 5870–9420 Hz. In particular, the guanine platelets exhibited a clear grid pattern at both 5870 and 9420 Hz (Figure 5A). Fast Fourier transfer (FFT) images showed ordered and isolated spots, indicating that the grid patterns were spatially uniform. The aqueous suspension of TiO_2_ particles exhibited a partially uniform grid pattern (Figure 5B, bottom panel) at 7600 Hz, whereas an amorphous pattern appeared at 7170 Hz, with a ring pattern in the FFT image (Figure 5B, top panel). In contrast, as shown in Figure 5C, the aqueous suspension of BaSO_4_ particles exhibited a diffraction pattern at 7230 Hz resembling that of the guanine platelet suspension at 1870 Hz.

**Figure 5.**
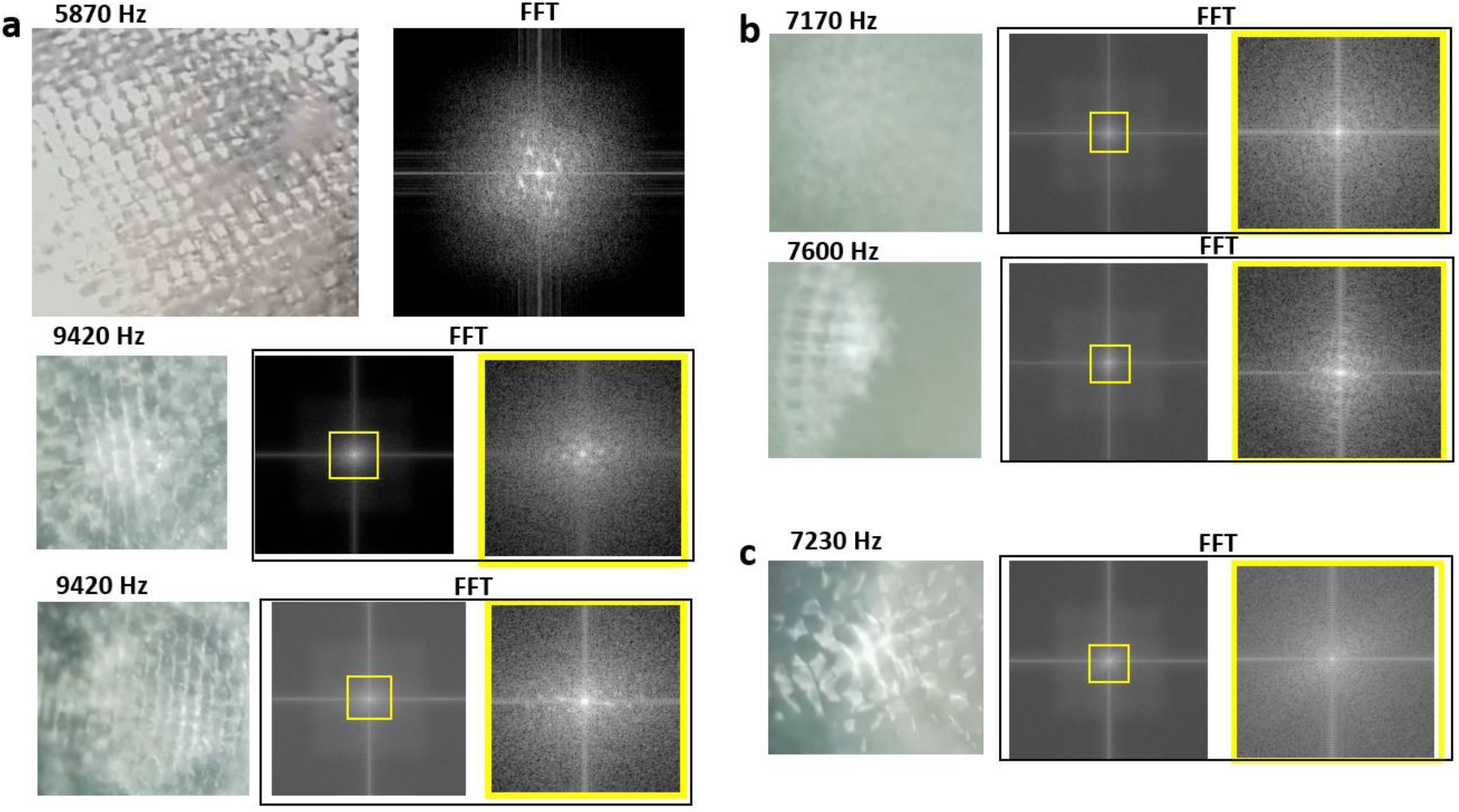
Comparison of standing lattice patterns of scattered light for water droplets containing three types of microparticles: (a) guanine platelets from fish, (b) TiO_2_ particles, and (c) BaSO_4_ particles. Representative patterns at frequencies of 5870–9420 Hz are shown for each material. The FFT pattern for each material is shown in the right (and center) panel(s) of each row. Right panels surrounded by a yellow box present a magnification of the center panel. The scale bar represents 200 μm.

These results suggest the presence of two types of light diffraction patterns on the water surface: locally appearing patterns (Figure 2, Figures 5B, C) and a grid pattern covering the entire surface of the water droplet (Figure 3, Figure 5A). The mechanism of these patterns is attributed to the diffusion of the water stream induced by the acoustic field transducer. As shown in Figure 2, a water stream was induced as the acoustic field frequency increased from 0 to 1870 Hz. The floating guanine platelets aligned along the stream, which enhanced the visualization of the flow (Figure 2a).

For the experimental conditions of density and volume for the water droplets studied herein, the guanine platelets exhibited a grid pattern of light diffraction at frequencies of 5870, 7420, and 9420 Hz, whereas no lattice pattern was observed at 1870 Hz. Figure 6C shows a summation of the intensity profiles along six horizontal lines. In the profiles, maximum peaks appeared every 100 μm along the water droplet containing guanine platelets. However, when the frequency of the acoustic field was shifted from 9420 to 9960 Hz, i.e., when less harmonic energy was transferred, the lattice pattern was slightly broken by a vortex flow (Figures 6D– F). A photograph of this droplet (Figure 6D) and two intensity profiles (Figure 6E, F) suggest the presence of diffraction peaks around the transmitted light (light reflected from the cover plate on which the water droplet was placed).

**Figure 6.**
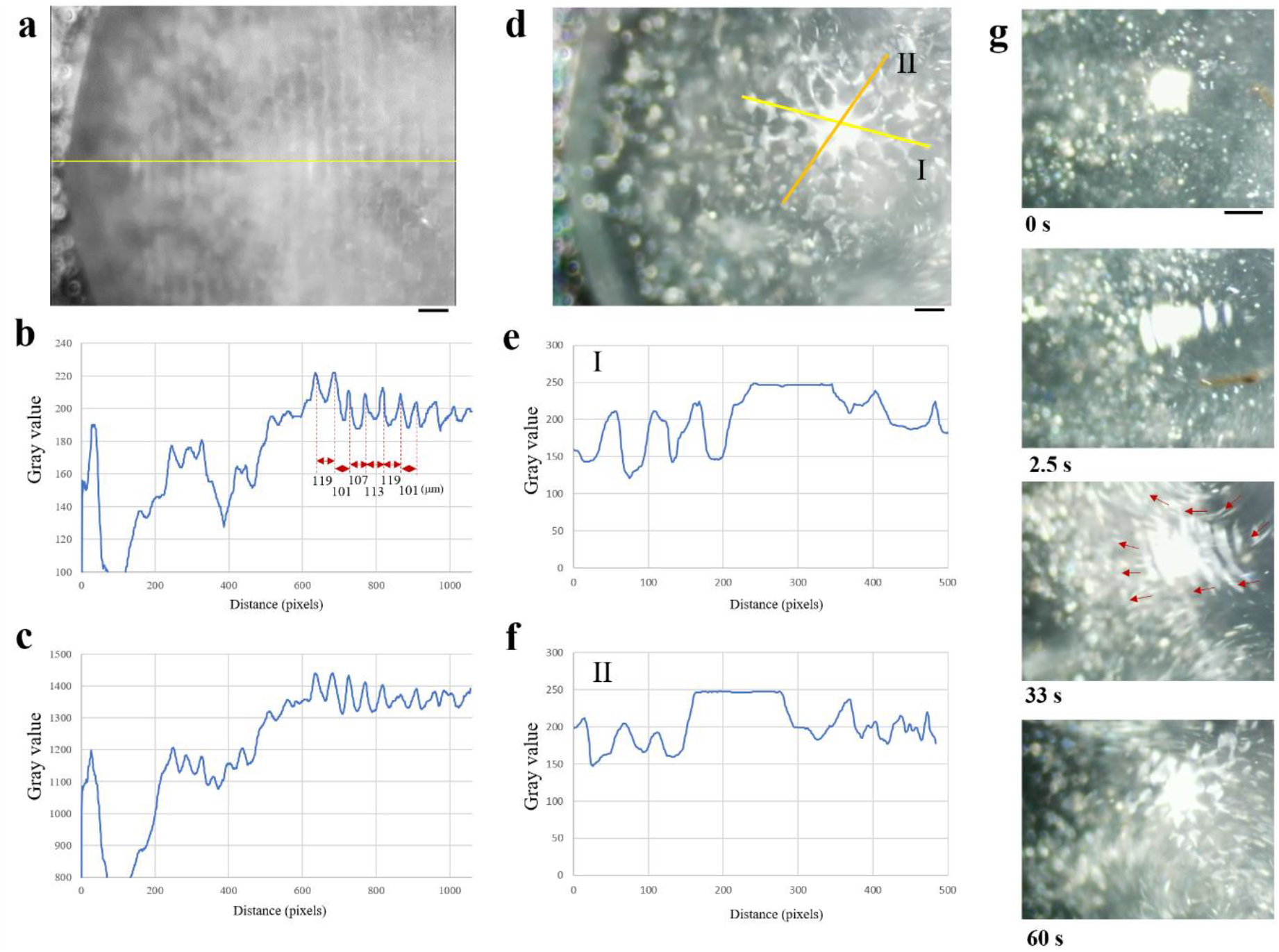
Analysis of the formation of standing lattice patterns of scattered light on water droplets containing guanine platelets from fish. (a) Grid of the diffraction pattern for acoustic field stimulation at 9420 Hz. The scale bar represents 200 μm. (b) Profile of intensity to gray along the horizontal line (yellow line) for the image shown in (a). (c) Summation of the intensity profiles along six horizontal lines parallel to the yellow line in (a). (d) Locally formed diffraction patterns for acoustic field stimulation at 9960 Hz. The scale bar represents 200 μm. (e) Profile of intensity to gray along line I shown in (d). (f) Profile of intensity to gray along line II shown in (d). (g) Growth of the diffraction pattern for acoustic field stimulation at 10000 Hz. The scale bar represents 250 μm.

A growing diffraction pattern was observed when the acoustic field frequency was shifted from 9420 to 10000 Hz, as shown in Figure 6G. A duration of several seconds was needed for the applied acoustic field with a rectangular wave to produce a one-dimensional lattice pattern. Within 30–60 seconds, a two-dimensional lattice pattern appeared. Thus, a sufficient amount of time was required for the lattice pattern to form on the water droplet, which most likely corresponds to the time needed to induce vortex flows in the particle suspension. Thereafter, the self-organized vortex flows containing particles provided a condition in which a periodical distribution of a distinct refractive index can induce a standing lattice pattern of scattered light.

## CONCLUSIONS

In this study, the effects of acoustic field stimulation on the optical surface image of a water droplet were observed. LED light was provided from the side onto a water droplet containing floating particles above a piezoelectric membrane plate. At a specific frequency of the applied acoustic field, a lattice or cellular pattern appeared. The lattice pattern maintained its position, overlapping with the background and the stream of water.

Compared with TiO_2_ and BaSO_4_ particles, biogenic guanine platelets from fish exhibited a clearer lattice pattern of reflected light over the surface of the water droplet. As the acoustic field frequency increased from 1900 to 10000 Hz, the surface lattice structure generated by the reflected light became more distinct.

Analyses of light intensity profiles of the lattice pattern of reflected light on the water droplet suggested the presence of diffraction peaks around the light transmitted through the guanine platelets in the water stream. The growth of a light diffraction pattern was also observed in the stream of platelets.

## AUTHOR INFORMATION

### Author Contributions

M.I. performed the experimental design, experiments, measurements and analyses. All part of manuscript and illustrations were prepared by M. I.

### Notes

The authors declare no competing financial interest.

## ACKNOWLEDGMENTS

This work was supported by JST-CREST “Advanced core technology for creation and practical utilization of innovative properties and functions based upon optics and photonics (Grant number: JPMJCR16N1).”

